# Cyclodextrin-Based Delivery of the Annexin A1 Mimetic Peptide Ac2-26 Enhances Anti-Inflammatory Effects and Prevents Dengue-Induced Lethality in Combination with Antiviral Therapy

**DOI:** 10.1101/2025.09.02.673789

**Authors:** Jenniffer Ramos Martins, Viviane Lima Batista, Laís Gomes Ramos, Celso Martins Queiroz-Junior, Talita Cristina Martins da Fonseca, Angélica Samer Lallo Dias, Leticia Soldati Silva, Felipe Rocha da Silva Santos, Ana Luiza de Castro Santos, Ivana Silva Lula, Felipe Emanuel Oliveira Rocha, Jéssica Aparecida Barsalini Pereira, Edvaldo Souza Lara, Thiago Moreno L. Souza, Lirlândia Pires Sousa, Mauro Martins Teixeira, Pedro Pires Goulart Guimarães, Vivian Vasconcelos Costa

## Abstract

Severe dengue is characterized by systemic inflammation, cytokine storm, vascular leakage, and hemorrhagic manifestations, largely driven by the host immune response to dengue virus (DENV) infection. Despite its burden, no licensed antivirals or host-directed therapies are currently available. Our group has previously identified Annexin A1 (AnxA1) as an endogenous regulator of inflammation in dengue. Treatment with the AnxA1 peptidomimetic, Ac_2-26_, improved clinical outcomes in murine models of severe dengue by promoting resolution of inflammation without affecting viral control. To explore new delivery strategies, we developed a novel formulation of Ac_2-26_ complexed with hydroxypropyl-β-cyclodextrin (CDX-Ac_2-26_). In DENV-2-infected A129 mice, both intraperitoneal and oral CDX-Ac_2-26_ improved clinical scores and reversed thrombocytopenia. Notably, CDX-Ac_2-26_ reduced mast cell degranulation, MCPT-1 plasma levels, and CCL2 expression in spleen, with no effect on viral titers, indicating a host-targeted mechanism and overcoming the anti-inflammatory effects of the free peptide. Intraperitoneal administration achieved the same efficacy as oral dosing with only one-third of the dose. Importantly, the combination of CDX-Ac_2-26_ with the antiviral nucleotide analog sofosbuvir fully prevented disease and mortality in infected mice, highlighting a combinatorial effect between host-directed and antiviral therapies. These findings underscore the therapeutic potential of anti-inflammatory/pro-resolving strategies in severe dengue and support the development of CDX-Ac_2-26_ as a novel adjunctive treatment. Combining anti-inflammatory and antiviral approaches may enhance efficacy and reduce treatment-associated toxicity, offering a promising path for clinical translation.

**What is already known on this topic?:**

- Plasma levels of Annexin A1 are inversely correlated with the severity of clinical outcomes in dengue virus infection.
- The Annexin A1 peptidomimetic Ac_2-26_ exhibits anti-inflammatory and pro-resolving effects against severe dengue in a murine model.

**What does this study add?:**

- The Ac_2-26_-cyclodextrin complex enhances the anti-inflammatory effects of the peptide against dengue virus infection, allowing for lower dosing and oral administration.
- The administration of the CDX-Ac_2-26_ with a nucleoside analog antiviral exhibits a combinatorial effectt, providing complete protection against the lethal outcome of severe dengue.

**What is the clinical significance?:**

- Current dengue treatment relies on symptomatic management, as no directed anti-inflammatory agents or antivirals are currently available. We have identified a novel host-targeted strategy to resolve the disease.
- Our findings strongly support CDX-Ac_2-26_ as a promising adjunctive treatment strategy in combination with antiviral therapy for severe dengue, highlighting the potential of combinatorial approaches.

## 1. INTRODUCTION

Dengue is an endemic viral disease that can present asymptomatically, with or without warning signs, and may progress to the most severe form of the disease (Ministério da Saúde., 2025). Annually, approximately 390 million individuals are infected with the dengue virus (DENV) worldwide, with at least 1% of infections progressing to severe dengue (SD). Although the proportion of severe cases is relatively low, the severity and potential for fatal outcomes must be considered, as the mortality rate of SD reaches approximately 20% (Bhatt et al., 2013). The progression to severe forms of the disease is associated with multiple factors, including age and prior infections with different DENV serotypes, which can trigger antibody-dependent enhancement (ADE) (Annsley et al., 2024), or the reactivation of heterologous memory T cells, a phenomenon known as original antigenic sin (Nikin-Beers el al., 2018). Moreover, viral factors such as serotype and genotype, and viral load, or even preexisting comorbidities, such as diabetes and cardiovascular diseases are also key determinants of an exacerbated inflammatory response which drives SD (Pulock et al., 2025).

After dengue virus inoculation into the host’s epidermis via the bite of an infected mosquito from the *Aedes* genus, infectious viral particles are capable of infecting resident dermal cells and disseminating through the lymphatic vessels into the circulatory system (McCracken et al., 2014). Viral recognition mechanisms trigger intracellular signaling cascades that culminate in the activation of transcription factors such as interferon regulatory factors (IRFs) and nuclear factor kappa B (NF-κB), by first line of immune defense cells, including Langerhans cells, monocytes, macrophages, endothelial cells, and mast cells. Located near blood vessels, mast cells contribute significantly to disease severity through the release of vasoactive mediators such as histamine, tryptase, and chymase (St John et al., 2013).

Following viral recognition, a local and systemic inflammatory response is initiated, leading to the recruitment of immune effector cells (eg: monocytes, macrophages, neutrophils, and lymphocytes) and the production of inflammatory mediators (eg: IL-1β, IL-10, IL-6, IL-8, IFN-γ and CCL2) aimed at containing pathogen dissemination (Bozza et al., 2008). However, if not correctly regulated, this immune response can become exacerbated, resulting in a cytokine storm an event closely associated with the pathophysiology of SD. This dysregulated process contributes to plasma leakage, thrombocytopenia, and tissue damage, reflecting a failure in the proper resolution of inflammation (Masyeni et al., 2024). In line with that, several research groups have focused on investigating the role of different pro-resolving mediators as potential therapeutic agents for the treatment of inflammatory conditions, both infectious and non-infectious (Costa et al., 2024). Among them, Annexin A1 (AnxA1) stands out as a protein with well-documented anti-inflammatory and pro-resolving properties.

AnxA1 is endogenously synthesized and widely expressed in different cell types, including leukocytes, lymphocytes, epithelial cells, and endothelial cells. It is recognized by the formyl peptide receptor (FPR) family, particularly FPR2/ALX (the lipoxin A4 receptor) (Sugimoto et al., 2019) acting as a potent anti-inflammatory and pro-resolving mediator exerting its effects by inhibiting leukocyte adhesion and migration inducing neutrophil apoptosis followed by clearance by macrophages, and promoting the differentiation of monocytes into macrophages with a pro-resolving phenotype (Perretti et al., 2023). In the context of DENV-induced inflammation our research group has demonstrated the relevance of AnxA1 as a potential anti-inflammatory and pro-resolving molecule. The results identified AnxA1 as a possible biomarker of disease severity, as plasma levels of this protein were reduced in infected patients compared to healthy individuals. Moreover, patients diagnosed with SD exhibited even lower levels of AnxA1 compared to those with mild clinical forms of the disease. In the same study, a murine model of DENV infection was employed, in which AnxA1-deficient (AnxA1^□/□^) mice developed more severe manifestations of the disease in comparison to wild-type (WT) animals, while administration of the mimetic AnxA1 peptide, Ac_2-26_ to infected animals significantly improved the clinical outcome, by promoting the resolution of DENV-induced inflammation (Costa et al., 2022).

The use of a peptidomimetic capable of reproducing the main functions of the full-length protein is of great interest (Yu et al., 2019). Thus, the development of optimized formulations represents a promising strategy to enable the systemic administration of such as Ac_2-26_. Cyclodextrins (CDs), a family of cyclic oligosaccharides composed of α-1,4-D-glucopyranoside units, can play an important role as carrier molecules for peptide delivery (Tokihiro et al., 2000). In the pharmaceutical context, CDs are particularly valued for their high biocompatibility, low toxicity, resistance to degradation, good aqueous solubility, and lack of immunogenicity features that make them excellent drug delivery vehicles (Ma et al., 2012).

In the present study, we aimed to develop a targeted therapeutic strategy for SD using a CD-based formulation to enhance the delivery of the Ac_2-26_ peptide. Our data demonstrated that the CDX-Ac_2-26_ formulation improved clinical outcomes in experimental SD, even at reduced doses or when administered orally demonstrating a selective anti-inflammatory effect, as the virus titers remained unaltered. Combining CDX-Ac_2-26_ with the antiviral Sofosbuvir, a nucleoside analog (Medina et al., 2023), has shown to be endowed with anti-flavivirus activity further improved the therapeutic effect and led to full survival of infected mice, highlighting the value of combined treatment strategies.

## 2. METHODS

### 2.1 Virus

The dengue virus strain used in this experiment was DENV-2 (Strain 3295: GenBank EU081177.1), kindly provided by Prof. Eng Eong Ooi from the Duke-NUS Medical School, Singapore. Viral stocks were propagated in VERO CCL-81 cells (kidney-derived cells from African green monkey *Cercopithecus aethiops*, BCRJ code 0343), maintained in a humidified incubator at 37 °C, using RPMI 1X culture medium supplemented with 1% antibiotics, 1% L-glutamine, 1% non-essential amino acids, and 10% fetal bovine serum (FBS), for 3-5 days. After propagation, the supernatant was collected and centrifuged at 600 × *g* for 10 minutes to remove cellular debris. The clarified supernatant was then subjected to viral concentration using a Vivacell 100 centrifugal concentrator (Sartorius, Germany), (2.000 × *g* for 10 minutes). The viral concentrate was aliquoted and stored at –80 °C. Viral titers were determined by plaque assay in VERO cells, with a final titer of 5.7 × 10C plaque-forming units per milliliter (PFU/mL).

### 2.2 Cytotoxicity and antiviral effect of cyclodextrins (CDs)

VERO CCL-81 cells, were cultured in 96-well plates, seeded at a density of 1 × 10C cells/well in RPMI 1X medium supplemented with 10% FBS and 1% antibiotics. After reaching >90% confluence, the cells were used for the assays described below. For cytotoxicity assessment, cells were exposed to four different cyclodextrins: β-Cyclodextrin, Sulfobutyl-ether-β-Cyclodextrin, Hydroxypropyl-β-Cyclodextrin, and Quaternary Ammonium-β-Cyclodextrin, at decreasing concentrations of 40 mM, 20 mM, 10 mM and 5 mM. The exposure lasted for 1 hour at 37 °C and 5% COC. Subsequently, the cells were washed with sterile 1X PBS and maintained in culture for 48 hours. Cell viability was assessed using the MTT assay (3-(4,5-dimethylthiazol-2-yl)-2,5-diphenyl tetrazolium bromide), according to the manufacturer’s instructions. To evaluate the antiviral effect, DENV-2 was incubated with the same cyclodextrins at the same concentrations (40 mM, 20 mM, 10 mM and 5 mM) for 1 hour at 37 °C in a 5% COC incubator, forming a single solution. This DENV-2 + CD solution was then added to VERO CCL-81 cells and incubated again for 1 hour under the same conditions. After incubation, the cells were washed and maintained in RPMI 1X medium supplemented with 2% FBS and 1% antibiotics for 48 hours. Cell viability was again assessed using the MTT assay. Supernatants were collected and stored at –80 °C for subsequent viral titer quantification.

### 2.3 Viral Titer Determination

Viral titration was performed on viral stocks, cell culture supernatants, and plasma and spleen samples collected from DENV-infected mice. Viral load was quantified by plaque assay using VERO CCL-81 cells, as previously described by (Costa et al., 2022). Initially, VERO CCL-81 cells were seeded into 24-well plates at a density of 1 × 10C cells per well. When the wells reached ∼90% confluence forming a uniform monolayer., viral stocks and plasma samples from infected mice were serially diluted in RPMI medium. Spleen samples were weighed, macerated using sterile porcelain mortars and pestles, and serially diluted in RPMI (supplemented with antibiotics and devoid of FBS) at a 10% weight/volume ratio. For each well, 100 μL of the respective sample dilutions were added: tissue samples (dilutions 10C² to 10CC), plasma (10C² to 10CC), or viral stocks (10C² to 10CC). A negative cell control was included by adding RPMI medium without virus to one well. Plates were incubated at 37 °C for 1 hour to allow viral adsorption, with gentle agitation every 15 minutes. After adsorption, the medium was removed, and the wells were washed with RPMI 1X. Subsequently, RPMI 1X medium containing 1.6% carboxymethylcellulose, antibiotics, and 2% FBS was added to each well. Plates were incubated for 3 days at 37 °C, during which cytopathic effects (CPE) were monitored using an inverted microscope. After the incubation period, the plates were fixed with 10% buffered formalin for at least 30 minutes and stained with 1% (w/v) crystal violet solution in deionized water. Plaques were then counted, and viral titers were calculated and expressed as PFU/mL.

### 2.4 Formulation of CDX-Ac_2-26_

The formulation of the Ac_2-26_ peptide in Hydroxypropyl-β-cyclodextrin was initially based on calculating the molecular weight of both the peptide (3089.46) and the cyclodextrin (1375.36), ensuring a homogenization at a concentration ratio of 1:10, respectively. After diluting the Ac_2-26_ peptide along with Hydroxypropyl-β-cyclodextrin in PBS 1X, lyophilization was carried out, followed by storage at -20°C.

### 2.5 NMR and HR-DOSY Methodology

NMR experiments were carried out on a Bruker AVANCE-NEO 600MHz spectrometer operating at a resonance frequency of 600.16 MHz, at 25 °C, using 5mm automatic probe TBI 600-S3 with z-grad. The compounds (5.0 mg each) were dissolved in a mixture of 90% H2O and 10% D2O (total volume of 0.6 mL), and High-Resolution Diffusion-Ordered Spectroscopy (HR-DOSY) and Saturation Transfer Difference (STD) NMR experiments were conducted to demonstrate the occurrence of complexation. ¹H NMR spectra were acquired using the WATERGATE5 technique to suppress the residual water signal. HR-DOSY experiments were performed using the stimulated echo sequence with bipolar gradient pulses, modified to include water suppression by excitation sculpting with gradients (STEBPESGP1S). The magnetic field pulse gradient duration (δ)1 and diffusion time (Δ) were optimized for each sample to achieve complete signal dephasing at the maximum gradient strength. A series of 16 STEBPESGP1S spectra with 16k data points were acquired for each NMR experiment. The δ and Δ values were 2.5 ms and 100 ms, respectively. The gradient pulses were incremented linearly from 2% to 98% of the maximum gradient strength. STD NMR spectra were acquired using the pulse sequence stddiffesgp.3. The experiments were carried out with the following parameters: a 90° pulse length (P1) of 10 µs, a relaxation delay (D1) of 2 s, 24 scans (NS), 16k data points (TD), and 4 dummy scans (DS). Selective saturation of the receptor was applied using Gaussian-shaped pulses (P11 = 80 ms) for a total saturation time (D20) of 2 s. Three irradiation frequencies were used: - 40.00 ppm (off-resonance), and 10.03 ppm and 0.75 ppm (on-resonance). STD spectra were obtained by subtracting the on- and off-resonance spectra using the standard stdsplit routine. All spectra were processed and analyzed using Bruker software TopSpin® version 4.1.4.

### 2.6 Animals

A129 mice, deficient for the gene encoding type I interferon receptor (IFN-α/β) (male and female), approximately 25 g, 8 weeks of age, were obtained from the vivarium of the University of São Paulo (USP) and housed in the vivarium of the Department of Biochemistry and Immunology of the Institute of Biological Sciences of the Federal University of Minas Gerais. The total number of animals used in this study was 210 mice. All mice were kept under specific pathogen-free conditions with controlled temperature and humidity (21 ± 2 °C/45%), light (12:12 h dark cycle) and with free access to pelleted/autoclavable chow and water. The animals were organized in microisolator boxes which were identified according to the experimental group. To promote environmental enrichment in animal care, pre-sterilized cardboard cylinders were inserted into the cages throughout the experimental period. All experimental procedures of this study were approved by the Animal Ethics Committee of the Federal University of Minas Gerais, Brazil, under protocol number 26/2023. Animal studies are reported in accordance with the ARRIVE guidelines (Percie du Sert et al., 2020) and the recommendations of the British Journal of Pharmacology (Lilley et al., 2020). There were no exclusions of animals.

### 2.7 Mouse infection

Mice were infected via the intraplantar (subcutaneous) route with an inoculum of 2×10C PFU/mouse diluted in a final volume of 30 μL. Control animals were injected with 30 μL of saline and comprised the no infected group. Following infection, clinical signs of disease were monitored daily. Mice that reached a 20% loss in body weight were immediately euthanized in accordance with humane endpoint criteria, following established ethical guidelines to minimize suffering.

### 2.8 Pharmacological treatments

#### 2.8.1 Dose-response intraperitoneal treatment with free peptide Ac_2-26_ in a systemic dengue model

Male and female mice aged eight weeks were subcutaneously in the paw inoculated with 2 × 10C PFU of DENV-2 in a volume of 30 μL. At 36 hours post-infection (p.i.), treatment with the free Ac_2-26_ peptide (GenScript) was administered intraperitoneally (100 μL) at three different doses: 6, 2 or 0,6 mg/kg. Subsequent treatments were given at 48 and 60 hours p.i. At 72 hours p.i., the animals were euthanized, and target tissues were collected for analysis. Throughout the experiment, clinical scores were monitored daily to assess disease progression.

#### 2.8.2 Comparison of the effect of treatment with free peptide Ac_2-26_ and formulated peptide CDX-Ac_2-26_ in a systemic dengue model

Eight-week-old male and female mice received a subcutaneous in the paw injection of DENV-2 at a dose of 2 × 10C PFU (30 μL). At 36 hours post-infection (p.i.), treatment was administered intraperitoneally with either Ac_2-26_ or CDX-Ac_2-26_ (100 μL) at a dose of 6mg/kg. Additional doses were given at 48 and 60 hours p.i. At 72 hours p.i., the animals were euthanized, and tissues of interest were collected for analysis. Throughout the experiment, clinical scores were monitored daily to assess disease progression.

#### 2.8.3 Comparison of the effects of CDX-Ac_2-26_ treatment in different administration routes in a systemic dengue model

Eight-week-old male and female mice were infected subcutaneously in the paw with an inoculum of 2 × 10C (30 μL). At 36 hours post-infection (p.i.), treatment with CDX-Ac_2-26_ was administered either intraperitoneally or orally (100 μL) at a dose of 6mg/kg. Additional doses were given at 48 and 60 hours p.i. At 72 hours p.i., the animals were euthanized, and tissues of interest were collected for analysis. Throughout the experiment, clinical scores were monitored daily to assess disease progression.

#### 2.8.4 Comparison of different doses of CDX-Ac_2-26_ treatment by the intraperitoneal route of administration in a systemic dengue model

DENV-2 was administered subcutaneously in the paw to eight-week-old male and female mice at a dose of 2 × 10C PFU (30 μL). At 36 hours post-infection (p.i.), treatment with CDX-Ac_2-26_ was administered intraperitoneally (100 μL) at three different doses: 6, 2 or 0,6 mg/kg. Additional doses were given at 48 and 60 hours p.i. At 72 hours p.i., the animals were euthanized, and tissues of interest were collected for analysis. Throughout the experiment, clinical scores were monitored daily to assess disease progression.

#### 2.8.5 Comparison of different doses of CDX-Ac_2-26_ treatment by the oral route of administration in a systemic dengue model

As previously described, eight-week-old male and female mice were subcutaneously in the paw infected with 2 × 10C PFU of DENV-2 (30 μL). At 36 hours post-infection (p.i.), treatment with CDX-Ac_2-26_ was administered orally (100 μL) at three different doses: 6, 2 or 0,6 mg/kg. Additional doses were given at 48 and 60 hours p.i. At 72 hours p.i., the animals were euthanized, and tissues of interest were collected for analysis. Throughout the experiment, clinical scores were monitored daily to assess disease progression.

#### 2.8.6 Assessment of survival outcomes following combined CDX-Ac_2-26_ and sofosbuvir treatment in a systemic dengue model

Eight-week-old male and female mice were infected subcutaneously in the paw with an inoculum of 2 × 10C (30 μL). One-hour post-infection (p.i.), treatment with the antiviral Sofosbuvir was initiated via oral administration at a dose of 60 mg/kg. At 36 hours p.i., treatment with Ac_2-26_ or CDX-Ac_2-26_ was initiated via intraperitoneal (100 μL) injection at a dose of 6mg/kg. Each treatment was administered every 12 hours, totaling 10 doses per treatment. Mice were monitored daily for clinical scores until day 14 post-infection. Humane endpoints were defined by a reduction of 20% of the mouse body weight.

### 2.9 Clinical Scoring

In order to evaluate the effects of infection and therapeutic interventions in infected mice, the following clinical parameters were evaluated: body weight loss (monitored daily using a semi-analytical scale, the maximum being 20%, more than that the animals were immediately euthanized), diarrhea, ocular discharge, piloerection, hunched posture, and reduced locomotor activity. Weight loss was scored based on percentage loss as follows: 0 = no weight loss; 0,5 = up to 5%; 1 = 5-10%; 1,5 = 10-15%; and 2 = 15-20%. For the other parameters, scoring was based on severity: 0 = absent; 0,5 = minimal; 1 = moderate; 1,5 = marked; and 2 = severe. The maximum possible clinical score was 12.

### 2.10 Euthanasia, Sample Collection, and Hematological Analysis

For blood collection, animals were anesthetized using a solution containing 100 mg/kg of ketamine and 10 mg/kg of xylazine. A midline laparotomy was performed, followed by blood collection from the inferior vena cava using heparinized syringes. Subsequently, animals were euthanized by anesthetic overdose followed by cervical dislocation for the collection of target tissues. In lethality experiments, blood from mice was collected at a single time point by puncture of the submandibular vein, in a volume of 30 μL (one drop) for platelet counting. Blood samples were transferred to heparinized tubes for platelet quantification and subsequent serum separation for cytokine quantification. Platelets were manually counted using a Neubauer chamber after a 1:100 dilution in 1% ammonium oxalate. Results are expressed as the total number of platelets per 10³ μL of blood. Whole blood was centrifuged to separate and collect serum (2.000 g for 5 minutes at 4 °C). The serum was then aliquoted for ELISA assays (stored at -20 °C).

### 2.11 Cytokine Quantification by ELISA

To quantify inflammatory mediators such as mast cell protease-1 (MCPT-1) and chemokines (CCL2), spleen fragments were homogenized in PBS containing protease inhibitors (0.1 mM phenylmethylsulfonyl fluoride, 0.1 mM benzethonium chloride, 10 mM EDTA, 20 KI aprotinin A) and 0.05% Tween 20, in the ratio of 0.1 g of tissue per mL of solution, using a tissue homogenizer (Power Gen 125, Fisher Scientific, Pennsylvania, USA). The homogenates were centrifuged at 2.000x*g* for 10 minutes at 4 °C (Centrifuge BR4, Jouan, Winchester, VA, USA), and the supernatants were collected and stored in 2 mL microtubes (Eppendorf, Brazil) at –20 °C for subsequent analysis. Serum and tissue samples were analyzed at a 1:5 dilution in PBS containing 0.1% bovine serum albumin (BSA), as previously standardized in our laboratory. ELISA assays were performed using antibody kits (R&D Systems, USA), following the manufacturer’s instructions.

### 2.12 Histological Evaluation

After euthanasia, mouse plantar paw pad was fixed in 4% paraformaldehyde in PBS (PFA 4%) for 2 days and subsequently processed through a series of increasing concentrations of ethanol, xylene, and liquid paraffin. Once processed, the tissues were embedded in paraffin, sectioned using a microtome, and mounted onto histological slides. The sections were stained with Toluidine Blue, and the total number of mast cells, intact and degranulated, was counted based on the metachromatic staining of the vesicles as previously described (St John et al., 2013).

### 2.13 Statistical Analysis

The data and statistical analysis complied with the recommendations of the British Journal of Pharmacology on experimental design and analysis (Curtis et al., 2022). Studies were designed to generate groups of equal size, using randomization and blinded analysis. Study sample size was calculated based on previous studies to achieve a 80% power (α=0.05) (Costa et al., 2022). Statistical analysis was performed for data where each sample size was at least N = 6. Data were tested for normality using Shapiro-Wilk test, and statistical significance was determined using GraphPad Prism 8 software for MacOs (GraphPad Software Inc., CA, USA). All results are expressed as mean ± SEM. Data were analyzed by two-way or one-way ANOVA, followed by the Tukey’s post hoc test. When two groups were evaluated, unpaired Student’s t-test was used. A value of p<0.05 was considered statistically significant, and are presented in each figure.

### 2.14 Literature Search

The bibliographic research for relevant electronic articles was conducted throughout the course of this study. Searches were performed using the Medline/PubMed database (https://pubmed.ncbi.nlm.nih.gov/) and the CAPES Periodicals Portal (Coordination for the Improvement of Higher Education Personnel – CAPES/MEC) (https://www-periodicos-capes-govbr.ezl.periodicos.capes.gov.br/index.php). Original articles, review papers, and textbook chapters written in English were selected based on their relevance to the study topic.

## 3. RESULTS

### 3.1 *In vitro* safety profile of Hydroxypropyl-**β**-Cyclodextrin supports its use in peptide formulation

Our group previously identified AnxA1 as an endogenous regulator of inflammation in dengue, with its peptidomimetic, Ac_2-26_, shown to improve clinical outcomes in murine models by promoting resolution of inflammation without impacting viral replication (Costa et al., 2022). In this and other prior studies, the effects were evaluated using a single intraperitoneal dose of Ac_2-26_ (6mg/kg) (Machado et al., 2020; Lara et al., 2025). To optimize dosing, we performed a dose–response experiment (6, 2 or 0,6mg/kg, intraperitoneally) in a severe dengue infection model. Results confirmed that the 6mg/kg dose was the most effective in improving clinical **(Supplementary** Fig. 1B) and hematological parameters **(Supplementary** Fig. 1C). However, this dose did not reduce CCL2 expression in the spleen **(Supplementary** Fig. 1D). While plasma levels of MCPT-1 were decreased **(Supplementary** Fig. 1E), no significant effect was observed on mast cell degranulation **(Supplementary** Fig. 1F). Importantly, treatment with Ac_2-26_ at any dose did not alter viral titers **(Supplementary** Fig. 1G), reinforcing its selective host-targeted mechanism of action. We hypothesize that formulating Ac2-26 would enhance its therapeutic efficacy by attenuating the dengue-induced cytokine storm. Therefore, we aimed to develop a delivery system capable of amplifying the peptide’s biological effects in the host and enabling its use in combination with antiviral candidates.

To develop the formulation, we investigated various cyclodextrins, compounds widely used for their ability to enhance solubility and modulate release profiles (de Castro et al., 2023). Our goal was to select a cyclodextrin with minimal cytotoxicity in Vero CC81 cells and no antiviral activity against DENV2, ensuring that any observed effects were attributable solely to improved peptide bioavailability. First, four cyclodextrins were evaluated in uninfected cells at different concentrations. β-cyclodextrin (β-CD; **Supplementary figure 2A**) and hydroxypropyl-β-cyclodextrin (HP-β-CD; **Supplementary figure 2B**) exhibited low cytoxicity, with cell viability remaining around 80% even at their highest concentrations tested. In contrast, Sulfobutyl-ether-β-cyclodextrin (SBE-β-CD; **Supplementary figure 2C**) and Quaternary Ammonium-β-cyclodextrin (QA-β-CD; **Supplementary figure 2D**) exhibited higher toxicity, reducing cell viability to approximately 60% and 20%, respectively, at the highest concentration.

We reproduced the assay using higher concentrations of the two cyclodextrins that had previously demonstrated low toxicity. Uninfected cells were treated with 40 mM of each cyclodextrin, in both biological and experimental triplicates, and viability assay was assessed. Consistent with earlier results, neither β-CD nor HP-β-CD showed significant cytotoxic effects **(Figure 1B and Figure 1C)**. Next, we assessed the antiviral activity of these cyclodextrins by measuring viral titers in the supernatants of DENV-2-infected cells. β-CD showed a marked antiviral effect against DENV2 **(Figure 1D),** while HP-β-CD led to only a modest decrease in viral titer **(Figure 1E),** even at the highest concentration. These findings supported the selection of HP-β-CD for formulation development, and the Ac_2-26_ peptide was subsequently formulated using it as the delivery vehicle.

**Figure 1.**
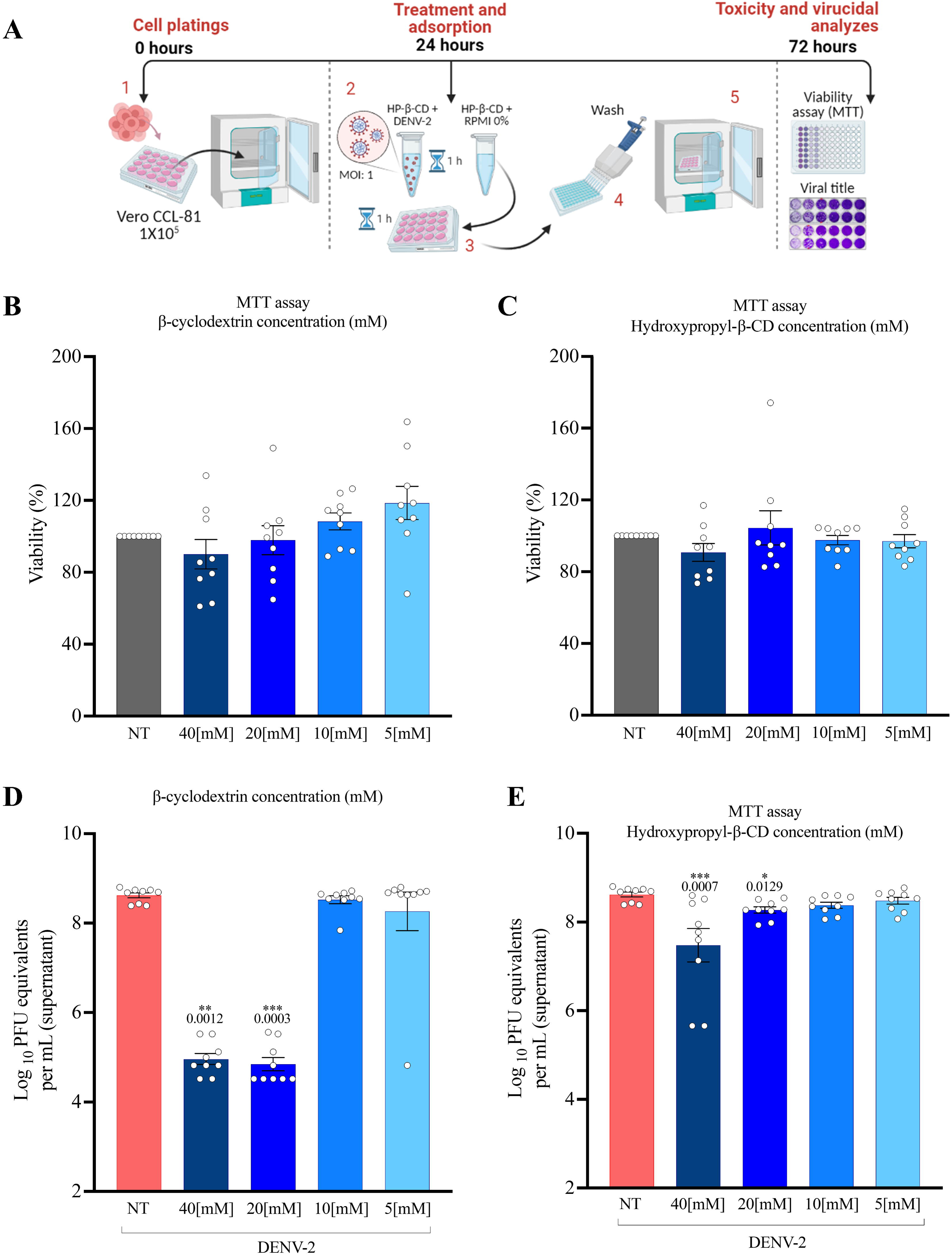
Evaluation of cytotoxicity and potential antiviral effect of β-cyclodextrin and hydroxypropyl-β-CD *in vitro*. Experimental design (A). Cytotoxicity assessment of the compounds was performed by evaluating NAD(P)H-dependent cellular oxidoreductase enzymes, which can reflect the number of viable cells present through plate reading (B) and (C). Plaque formations were counted, and sample titers were determined and expressed in PFU/mL (D) and (E). Results are expressed as mean ± SEM (biological and experimental triplicates N = 9 each group) and are represented with *p < 0.05, **p < 0.01, ***p < 0.001, and ****p < 0.0001 compared to the NT group. To assess the normality of the data distribution, the Shapiro-Wilk test was used. Parametric data were evaluated by ANOVA and Tukey’s post-test. Non-parametric data were subjected to non-parametric analysis of variance by the Kruskal-Wallis test followed by Dunn’s multiple comparisons post-test, with values considered statistically significant when p < 0.05.

### 3.2 NMR reveals specific interaction and complex formation between Ac_2-26_ and HP-**β**-CD

We formulated the complex (CDX-Ac_2-26_) and confirmed successful complex formation through physicochemical characterization. The results obtained from HR-DOSY experiment **(Figure 2B)** revealed differences in the translational diffusion coefficients among the individual components the Ac_2-26_ peptide (log(D) ≈ -9.65), HP-β-CD (log(D) ≈ -9.40), and the CDX-Ac_2-26_ complex (log(D) ≈ -9.00). These shifts indicate reduced molecular mobility consistent with complex formation. To further validate complex formation, STD NMR experiments were conducted. Irradiation frequencies were selected to excite the NH proton of tryptophan (10.03 ppm) and the internal methyl groups (0.75 ppm) of Ac2-26 **(Figure 2C)**. Upon irradiation at 0.75 ppm, saturation was transferred to the methyl hydrogens of Ac_2-26_, resulting in an observable STD signal at 1.08 ppm, corresponding to HP-β-CD. Additional signals in the 3.3–4.0 ppm region, characteristic of HP-β-CD, were also observed, indicating intermolecular interaction with the peptide **(Figure 2D)**. Similarly, irradiation at 10.03 ppm (NH of tryptophan) produced STD signals corresponding to HP-β-CD, suggesting that this aromatic region of Ac_2-26,_ including the tryptophan residue, is also involved in interactions with the cyclodextrin **(Figure 2E)**.

**Figure 2.**
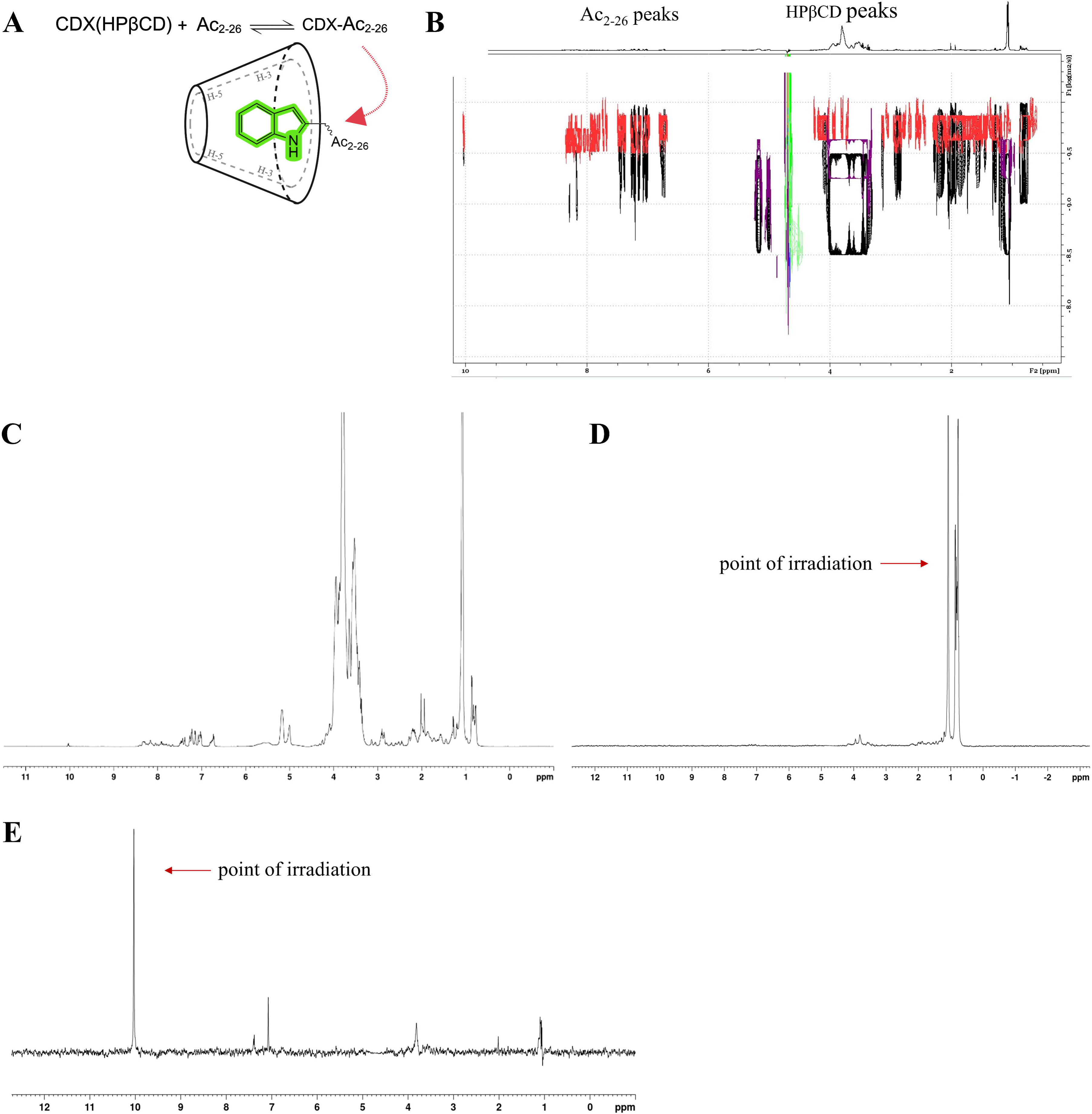
Characterization of the CDX-AC_2-26_ Complex by HR-DOSY and NMR Spectroscopy. Representative drawing of the interaction between Ac_2-26_ and HP-β-CD (A). HR-DOSY spectrum acquired in H2O/D2O (10%) at 600 MHz and 25°C, showing the diffusion coefficients of Ac_2-26_ (red), HP-β-CD (purple), and the CDX-AC_2-26_ complex (black), highlighting the different diffusion coefficients (B). ¹H NMR spectrum of the CDX-AC_2-26_ complex recorded in H2O/D2O (10%) at 600 MHz and 25°C (C); STD NMR spectrum upon irradiation at 0.75 ppm (D); STD NMR spectrum upon irradiation at 10.03 ppm (E), confirming the intermolecular interaction of the tryptophan residues of Ac_2-26_ and HP-β-CD.

### 3.3 CDX-Ac_2-26_ demonstrated improved effects compared to free Ac_2-26_, potentiating the reduction of pro-inflammatory mediators induced by DENV-2 infection

We then investigated whether the CDX-Ac_2-26_ would improve the effects when compared to the free Ac_2-26_ in a murine DENV2 infection model. A129 mice infected with DENV-2 were treated intraperitoneally with either the Ac_2-26_ or the CDX-Ac_2-26_ at a dose of 6mg/kg, 36 hours after infection, every 12 hours. We also included a group treated only with HP-β-CD (CDX) to verify any possible effect of the formulation excipient (**Figure 3A**). Both Ac_2-26_ treatments significantly improved the clinical score (**Figure 3B**) and prevented infection-induced thrombocytopenia (**Figure 3C**). However, only the CDX-Ac_2-26_ reduced the levels of the pro-inflammatory mediator CCL2 in the spleen (**Figure 3D**), and the plasma levels of MCPT-1 (**Figure 3E**), as well as the local mast cell degranulation in the hind paw plantar pad of A129 mice infected with DENV (**Figure 3F and Figure 3G**). Both treatments did not interfere with the host’s ability to control the infection, with similar viral titers observed in both groups (**Figure 3H**). Overall, the results demonstrate that the administration of the CDX-Ac_2-26_ enhanced its effects when compared to the free Ac_2-26_. The findings are important in demonstrating that peptides can have their effects enhanced when formulated in complexes.

**Figure 3.**
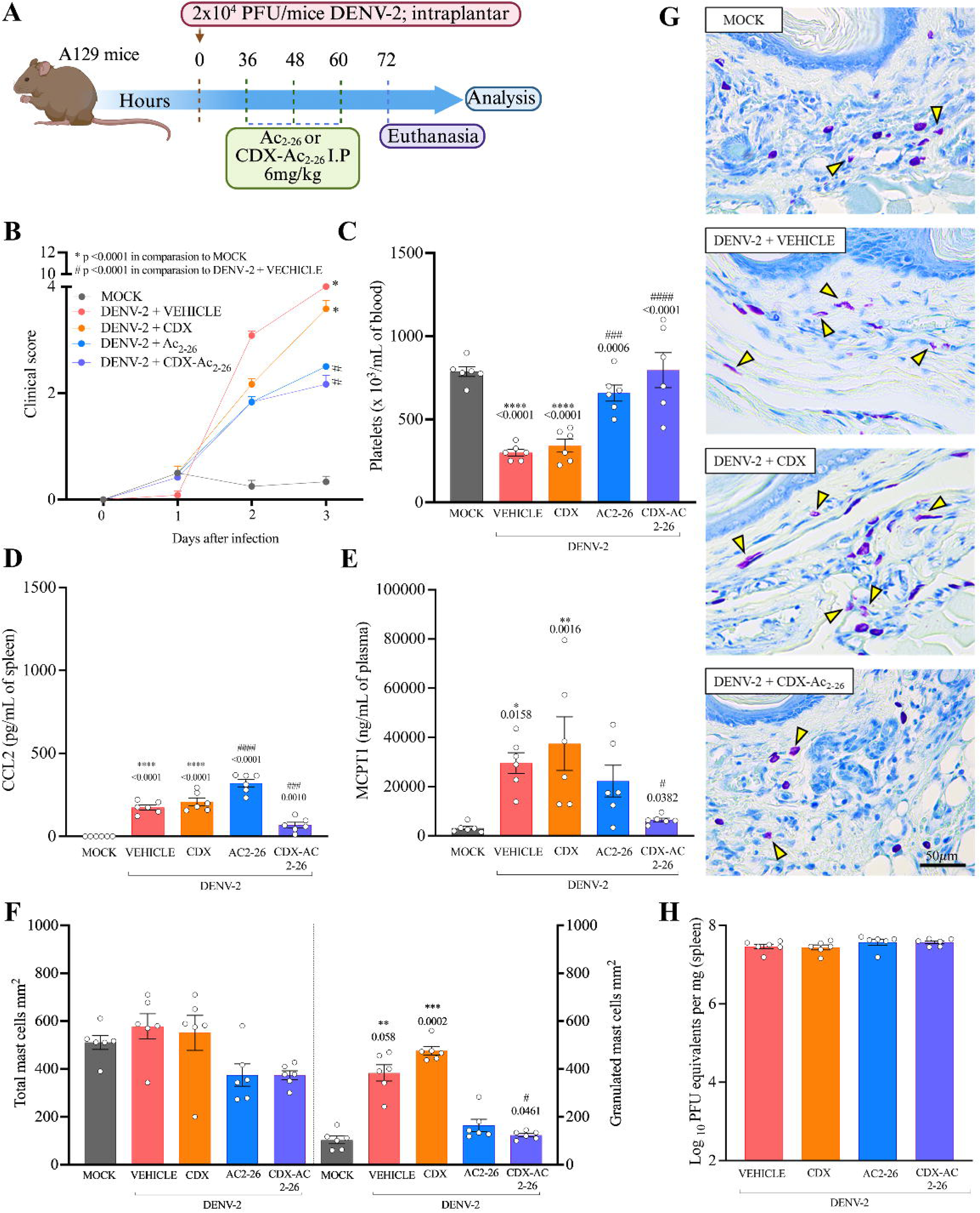
Comparison of the effect of treatment with free peptide Ac_2-26_ and formulated peptide CDX-Ac_2-26_ in a systemic dengue model. Experimental design (A). Evaluated clinical parameters (weight loss, diarrhea, piloerection, ocular secretion, low locomotor activity, and hunched posture) (B). Platelet count manually in a Neubauer chamber, with results presented as total platelet number/10³ μL of blood (C). Levels of inflammatory mediators measured by ELISA Kit, spleen CCL2 (D) and plasma levels of MCPT-1 (E). Manual counting of total mast cells, intact and degranulated locally in the paw (F) and representative images of histological sections of the plantar pad stained with toluidine blue dye (G). Viral titers in the spleen of infected mice were determined by plaque counting and sample titration, expressed in PFU/mL (H). Results are expressed as mean ± SEM (N = 6 animals in each group) and are represented with *p < 0.05, **p < 0.01, ***p < 0.001, and ****p < 0.0001 compared to the MOCK group. #p < 0.05, ##p < 0.01, ###p < 0.001, and ####p < 0.0001 compared to the DENV + VEHICLE group. To assess the normality of data distribution, the Shapiro-Wilk test was used. Parametric data were evaluated by ANOVA and Tukey’s post-test. Non-parametric data were subjected to non-parametric analysis of variance by the Kruskal-Wallis test followed by Dunn’s multiple comparisons post-test, with values considered statistically significant when p < 0.05. Outliers were not detected.

### 3.4 The therapeutic effects of CDX-Ac_2-26_ were preserved following oral administration

It has been highlighted the importance of administration routes, as drugs are generally better accepted by patients when delivered through less invasive such as the oral route. Based on this, we next investigated the effect of CDX-Ac_2-26_ given by oral route. To that end, A129 mice were treated with the CDX-Ac_2-26_ at a dose of 6mg/kg via intraperitoneal or oral route. Treatment via both routes resulted in an improvement in the clinical score and the treatment of uninfected mice no adverse events were observed (**Figure 4B**) and the prevention of the infection-induced thrombocytopenia (**Figure 4C**). Additionally, both administration routes reduced CCL2 levels in the spleen (**Figure 4D**), as well as plasma levels of MCPT-1 (**Figure 4E**), and were associated with decreased local mast cell degranulation in the footpad of A129 mice infected with DENV2 (**Figure 4F and Figure 4G).** Furthermore, neither treatment impaired the host’s ability to control the infection, as viral titers remained unchanged (**Figure 4H**). Overall, the results indicate that the CDX-Ac_2-26_ enables its administration through different routes, including intraperitoneal and oral delivery.

**Figure 4.**
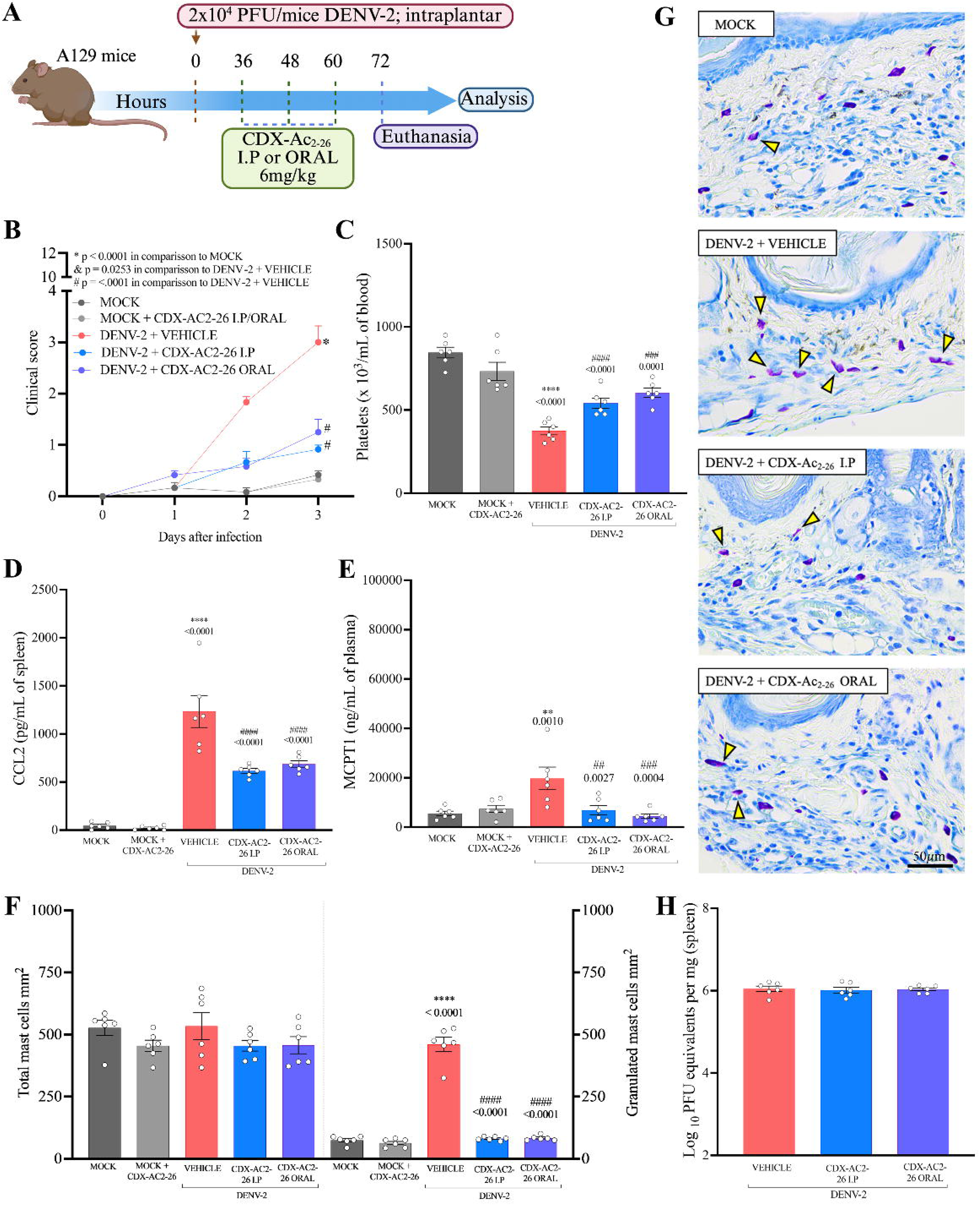
Comparison of the effects of CDX-Ac_2-26_ treatment in different administration routes in a systemic dengue model. Experimental design (A). Clinical parameters assessed (weight loss, diarrhea, fur bristling, ocular secretion, reduced locomotor activity, and hunched posture) (B). Manual platelet count using a Neubauer chamber, with results expressed as total platelet number/10³ μL of blood (C). Levels of inflammatory mediators measured by ELISA kit, CCL2 (D), and plasma levels of MCPT-1 (E). Manual counting of total, intact, and degranulated mast cells in the paw (F), and representative images of histological sections of the footpad stained with toluidine blue (G). Viral titers in the spleen of infected mice were determined by plaque formation assays, with results expressed in PFU/mL (H). Results are expressed as mean ± SEM (N = 6 animals in each group) and are represented with *p < 0.05, **p < 0.01, ***p < 0.001, and ****p < 0.0001 compared to the MOCK group. #p < 0.05, ##p < 0.01, ###p < 0.001, and ####p < 0.0001 compared to the DENV + VEHICLE group. The Shapiro-Wilk test was used to assess the normality of data distribution. Parametric data were analyzed using ANOVA followed by Tukey’s post-hoc test. Non-parametric data were analyzed using the Kruskal-Wallis test followed by Dunn’s multiple comparisons post-hoc test, with statistical significance considered for p < 0.05. Outliers were not detected.

### 3.5 CDX-Ac_2-26_ allows a threefold lower dose to be used via the intraperitoneal route while preserving efficacy

We then questioned whether treatment with the CDX-Ac_2-26_ would allow the peptide to be administered at a lower dose. For that, A129 mice were treated with the formulated peptide (CDX-Ac_2-26_) at doses of 6, 2 or 0,6mg/kg in intraperitoneal route (**Figure 5A**). Treatments at doses of 6 or 2mg/kg improved the clinical score (**Figure 5B**) and all doses prevented infection-induced thrombocytopenia even if partially (**Figure 5C**). Administration of 6 or 2mg/kg doses reduced CCL2 levels in the spleen (**Figure 5D**), decreased plasma levels of MCPT-1 (**Figure 5E**), and attenuated local mast cell degranulation in the footpad of A129 mice infected with DENV (**Figure 5F**). As we can see in the representative histological images of the skin (**Figure 5G**). Again, viral titers remained unchanged (**Figure 5H**). These results demonstrate that the CDX-AC_2-26_ enables the peptide to be administered a threefold lower dose (2mg/kg) while maintaining the same efficacy of the higher dose (6mg/kg).

**Figure 5.**
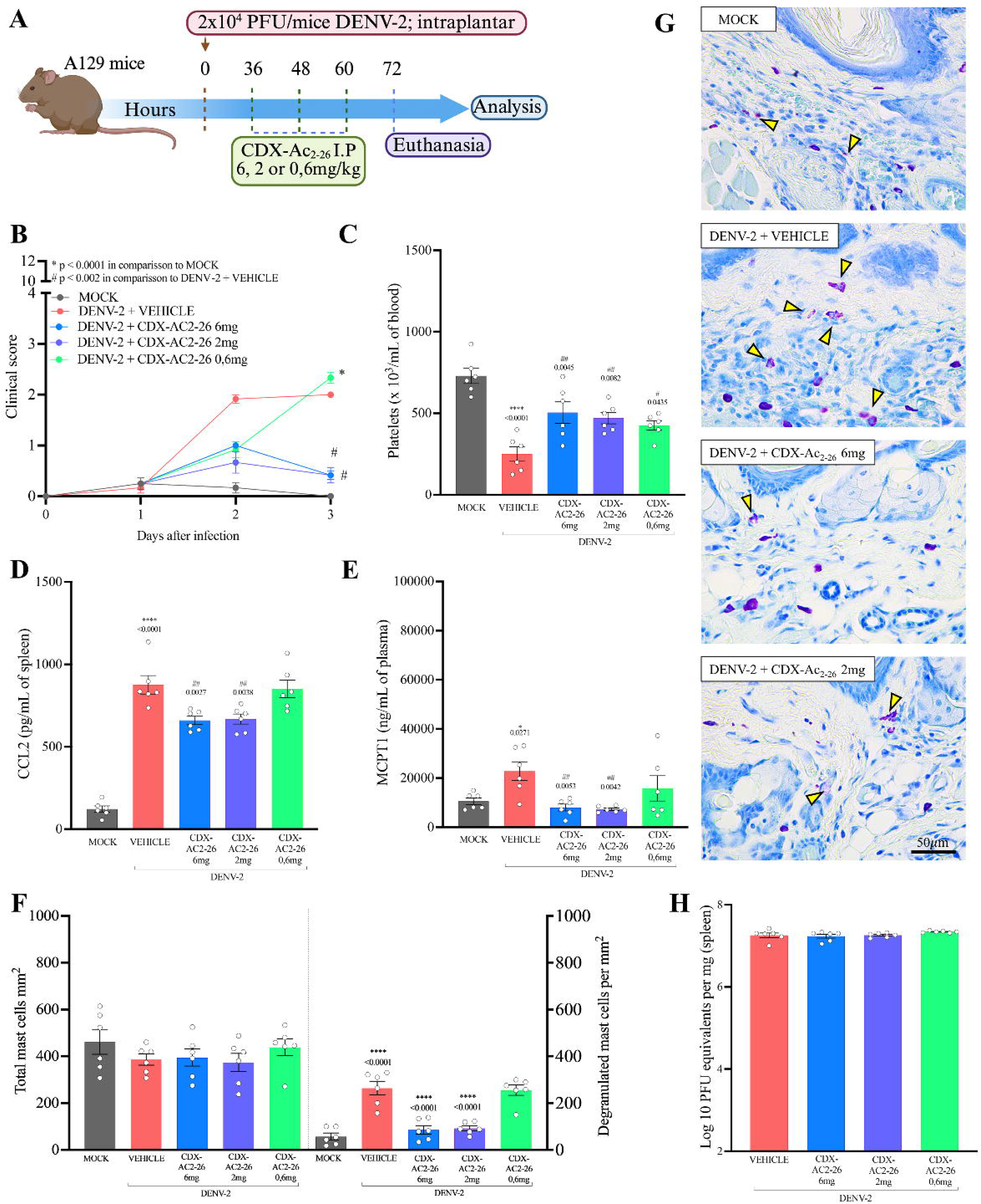
Comparison of different treatment doses of CDX-Ac_2-26_ by the intraperitoneal route of administration in a systemic dengue model. Experimental design (A). Clinical parameters assessed (weight loss, diarrhea, fur bristling, ocular secretion, reduced locomotor activity, and hunched posture) (B). Manual platelet count using a Neubauer chamber, with results expressed as total platelet number/10³ μL of blood (C). Levels of inflammatory mediators measured by ELISA kit, CCL2 (D), and plasma levels of MCPT-1 (E). Manual counting of total, intact, and degranulated mast cells in the paw (F), and representative images of histological sections of the footpad stained with toluidine blue (G). Viral titers in the spleen of infected mice were determined by plaque formation assays, with results expressed in PFU/mL (H). Results are expressed as mean ± SEM (N = 6 animals in each group) and are represented with *p < 0.05, **p < 0.01, ***p < 0.001, and ****p < 0.0001 compared to the MOCK group. #p < 0.05, ##p < 0.01, ###p < 0.001, and ####p < 0.0001 compared to the DENV + VEHICLE group. The Shapiro-Wilk test was used to assess the normality of data distribution. Parametric data were analyzed using ANOVA followed by Tukey’s post-hoc test. Non-parametric data were analyzed using the Kruskal-Wallis test followed by Dunn’s multiple comparisons post-hoc test, with statistical significance considered for p < 0.05. Outliers were not detected.

These results motivated us to extend the investigation to the oral route, emphasizing the importance of identifying the most effective route of administration while using the lowest possible therapeutic dose. In this scenario, treatment at the highest dose (6mg/kg) were more effective in reducing clinical scores (**Figure 6B**), and in preventing infection-induced thrombocytopenia, even if partially (**Figure 6C**). Administration in the two largest doses (6 and 2mg/kg) reduced CCL2 levels in the spleen (**Figure 6D**) and plasma levels of MCPT-1 in all doses (**Figure 6E**). Only the highest dose (6mg/kg) attenuated local mast cell degranulation in the footpad of A129 mice infected with DENV (**Figure 6F and 6G)**). As expected, none of the treatments interfered with viral titers, as they remained unchanged. (**Figure 6H**). All protective effects of CDX-AC_2-26_ were maintained only at the highest oral dose.

**Figure 6.**
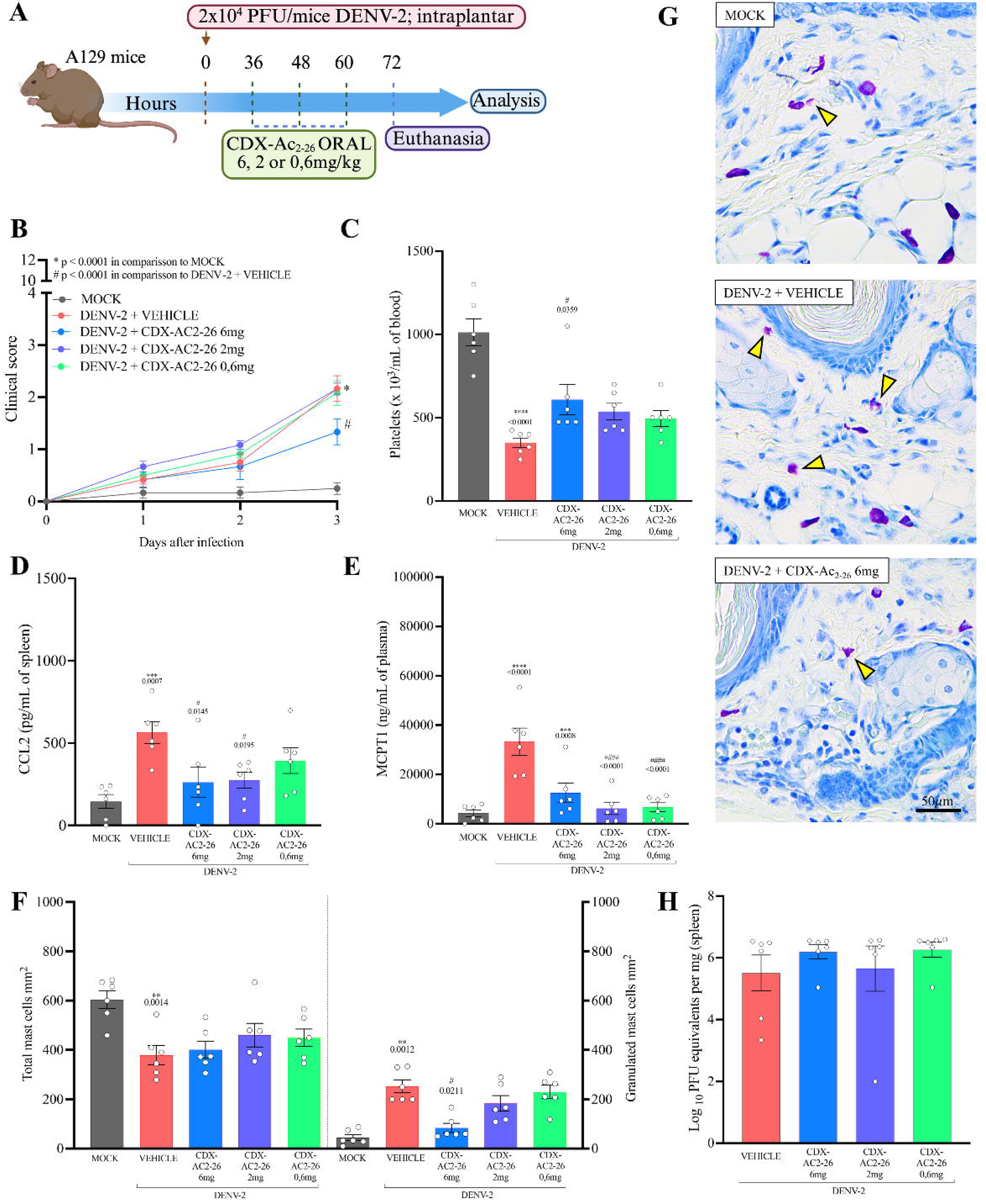
Comparison of different doses of CDX-Ac_2-26_ treatment by the oral route of administration in a systemic dengue model. Experimental design (A). Clinical parameters assessed (weight loss, diarrhea, fur bristling, ocular secretion, reduced locomotor activity, and hunched posture) (B). Manual platelet count using a Neubauer chamber, with results expressed as total platelet number/10³ μL of blood (C). Levels of inflammatory mediators measured by ELISA kit, CCL2 (D), and plasma levels of MCPT-1 (E). Manual counting of total, intact, and degranulated mast cells in the paw (F), and representative images of histological sections of the footpad stained with toluidine blue (G). Viral titers in the spleen of infected mice were determined by plaque formation assays, with results expressed in PFU/mL (H). Results are expressed as mean ± SEM (N = 6 animals in each group) and are represented with *p < 0.05, **p < 0.01, ***p < 0.001, and ****p < 0.0001 compared to the MOCK group. #p < 0.05, ##p < 0.01, ###p < 0.001, and ####p < 0.0001 compared to the DENV + VEHICLE group. The Shapiro-Wilk test was used to assess the normality of data distribution. Parametric data were analyzed using ANOVA followed by Tukey’s post-hoc test. Non-parametric data were analyzed using the Kruskal-Wallis test followed by Dunn’s multiple comparisons post-hoc test, with statistical significance considered for p < 0.05. Outliers were not detected.

### 3.6 Combination therapy with CDX-Ac_2-26_ and an antiviral agent prevents DENV-induced mortality in mice

CDX-Ac_2-26_ exhibited anti-inflammatory activity without impairing the host’s ability to control viral replication, as viral titers remained stable. Based on this, we tested whether treatment could improve survival, particularly when combined with an antiviral agent. As previously standardized, our 2×10C inoculum model effectively induces a range of inflammatory responses, such as cytokine production, but becomes highly lethal by day 7 due to its mimicry of severe infection. Therefore, to assess animal lethality within a broader yet still lethal therapeutic window, we used a lower inoculum of 2×10². All infected and untreated animals succumbed by day 9 post-infection, and those treated with Ac_2-26_ alone by day 11. Treatment with CDX-Ac_2-26_ alone resulted in a 33% survival rate, whereas Sofosbuvir alone increased survival to 50%. Remarkably, combined treatment with CDX-Ac_2-26_ and Sofosbuvir resulted in 100% survival **(Figure 7B)**. On day 14, surviving animals displayed clinical scores close to baseline (**Figure 7C)**. Treatment with Sofosbuvir did not prevent thrombocytopenia **(Figure 7D)**, but in an independent experiment performed under the same conditions, treatment with the antiviral reduced viral titers in the plasma and spleen of infected mice **(Figure 7E)**. CDX-Ac_2-26_ reduces inflammation while Sofosbuvir directly inhibits viral replication. This finding emphasizes the value of adjunctive strategies targeting SD.

**Figure 7.**
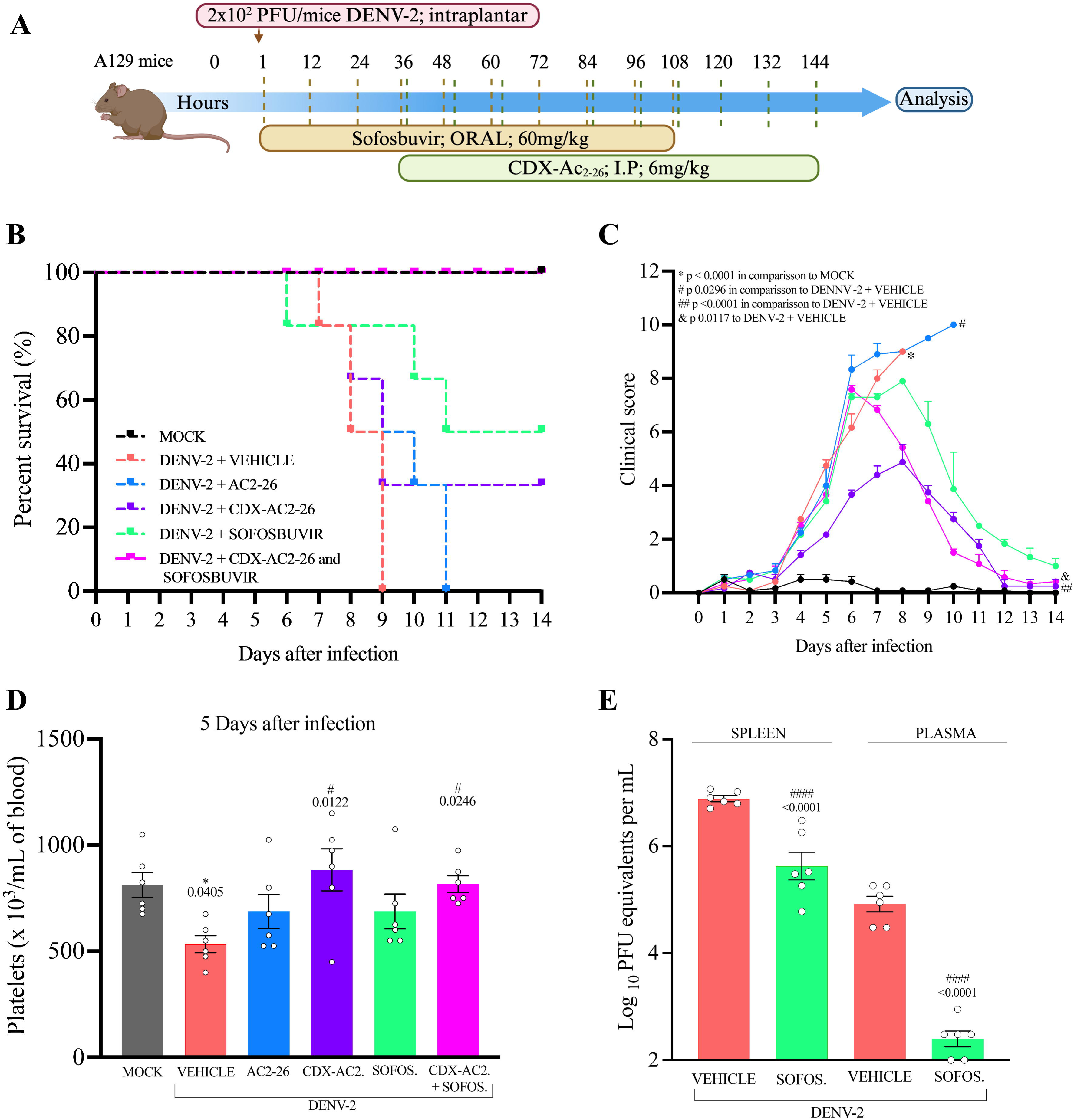
Effect of the combined treatment with CDX-Ac_2-26_ and Sofosbuvir in survival rates, clinical signs and viral load in a systemic dengue model. Experimental design (A). Percentage of survival (B). Clinical parameters evaluated (weight loss, diarrhea, ruffled fur, ocular discharge, reduced locomotor activity and hunched posture) (C). Blood collection on day 5 after infection by puncture of the submandibular vein and manual platelet count using a Neubauer chamber, with results expressed as total platelet number/10³ μL of blood (D). Viral titers in plasma and spleen of infected mice were determined by means of an independent experiment under the same conditions, by plaque formation assays, with results expressed in PFU/mL (E). Results are expressed as mean ± SEM (N = 6 animals in each group) and are represented with *p < 0.05, **p < 0.01, ***p < 0.001, and ****p < 0.0001 compared to the MOCK group. #p < 0.05, ##p < 0.01, ###p < 0.001, and ####p < 0.0001 compared to the DENV + VEHICLE group. The Shapiro-Wilk test was used to assess the normality of data distribution. Parametric data were analyzed using ANOVA followed by Tukey’s post-hoc test. Non-parametric data were analyzed using the Kruskal-Wallis test followed by Dunn’s multiple comparisons post-hoc test, with statistical significance considered for p < 0.05. Outliers were not detected.

## 4. DISCUSSION

Published results showed that reduced AnxA1 levels in DENV-infected individuals are associated with more severe clinical manifestations, underscoring its importance in modulating inflammation and the therapeutic potential of its peptidomimetic, Ac2_-26_, in attenuating inflammation in murine models of dengue (Costa et al., 2022). Here, we show that this effect is dose-dependent, but even at the highest dose of Ac_2-26_, not all clinical parameters were attenuated. This limitation may be attributed to the peptide’s low bioavailability, simple structure and lower *in vivo* stability compared to native AnxA1, which may compromise its efficacy (Broering et al., 2024; Qin et al., 2021). Additionally, high affinity for plasma proteins may reduce the peptide available to interact with target tissues, requiring higher doses to achieve therapeutic concentrations (Diao et al., 2013). Therefore, formulation strategies have emerged as a promising approach to overcome limitations associated with peptide administration (Mendes et al., 2024). We tested a formulation of the peptide in cyclodextrin, which demonstrated better attenuation of the clinical and inflammatory parameters induced by SD, even when administered in a dose three times lower and orally.

Among the carrier systems explored, CDs have gained prominence due to their broad applicability and safety profile in pharmacology. Their molecular architecture, with primary hydroxyl groups on the narrow edge and secondary hydroxyls on the wider edge, enables hydrogen bonding. CDs possess amphiphilic character, with a hydrophobic inner cavity and hydrophilic outer surface, making them well-suited for forming inclusion complexes with bioactive molecules (de Castro et al., 2023). Several CD-based pharmaceutical formulations are approved for clinical use, reinforcing their potential as effective carriers (Upadhyay et al., 2023). Due to the limited solubility and stability of Ac_2-26_, we developed a cyclodextrin-based formulation to improve its delivery and bioavailability. Among CDs, β-CDs were selected for their intermediate cavity size, compatible with bioactive molecules (de Paula et al., 2011). HP-β-CD, a derivative with enhanced solubility and reduced toxicity (Rajamohan et al., 2025), was the most candidate. Our *in vitro* studies demonstrated that HP-β-CD exhibited low cytotoxicity and minimal antiviral activity against DENV-2. Selecting a CD with negligible antiviral activity was to ensure that any observed therapeutic effects could be attributed to the peptide. This is particularly important given that some CDs, especially those with affinity for cholesterol, can sequester membrane lipids and disrupt viral envelopes, thereby exhibiting antiviral activity against enveloped viruses (Bezerra et al., 2022; Carro et al., 2013).

Based on these results, we developed a formulation CDX-Ac_2-26_ and evaluated interaction analyses to validate complex formation. Changes in the diffusion coefficient observed through HR-DOSY experiments suggest the presence of interactions between HP-β-CD and the Ac_2-26_ peptide. The results suggest the formation of a complex stabilized by weak interactions, such as hydrogen bonding and hydrophobic forces. The slight reduction in molecular mobility suggests an increased hydrodynamic radius, likely due to partial inclusion of the peptide within the cyclodextrin cavity (Walpole et al., 2019; Lula., 2011). Our findings support the hypothesis that both the internal methyl groups of HP-β-CD and the tryptophan residues in the Ac_2-26_ sequence contribute to the stabilization and formation of the inclusion complex.

We investigated CDX-Ac_2-26_ *in vivo* using A129 mice. This animal represents a robust tool for studying severe dengue as they are highly susceptible to viral infection due to deletion of genes encoding type IFN-α/β, resulting in a compromised innate antiviral response (Zandi et al., 2019; Santos et al., 2024). The use of wild type animals is limited because they are not considered natural reservoirs for dengue virus and are not susceptible to severe infection. We refined our model employing an intraplantar (subcutaneous) inoculation, which resembles the natural infection mechanism mediated by the mosquito bite (Mota et al., 2020). This led, on the third day post-infection, to high viremia and triggers an exacerbated inflammatory response, characterized by excessive production of inflammatory mediators. Treatment with Ac_2-26_ or CDX-Ac_2-26_ was initiated 36 hours after infection, aiming to preserve the host’s initial inflammatory response, crucial for controlling viral replication aiming to modulate excessive inflammation (Tavares et al., 2022). CDX-Ac_2-26_ and it demonstrated greater efficacy in attenuating the clinical and inflammatory parameters induced by DENV infection *in vivo* compared to the free peptide. These findings support the hypothesis that peptide complexation in CDs contributes to increased stability *in vivo*.

CDX-Ac_2-26_ proved effective in attenuating clinical and inflammatory parameters associated with DENV infection, including prevention of clinical score worsening and thrombocytopenia. This protection can be explained since AnxA1 has been described as a modulator of platelet function, reducing platelet activation, aggregate formation, and thrombosis (Vital et al., 2020). These effects can be partly mediated by inhibition of thrombin-induced αIIbβ3 integrin activation, suggesting an additional mechanism by which AnxA1 contributes to control of vascular complications associated with dengue (Senchenkova et al., 2019; Vital et al., 2016). In experimental study, AnxA1 inhibited expression of endothelial adhesion molecules such as E-selectin and VCAM-1 in Human Umbilical Vein Endothelial Cells, resulting in decreased adhesion of a human monocytic leukemia cell line to the endothelial monolayer (Zeng et al., 2025), which may also account for the reduced CCL2 levels observed with treatment the CDX-Ac_2-26_. AnxA1 also exerts anti-inflammatory and pro-resolving effects by inhibiting leukocyte adhesion and migration, inducing neutrophil apoptosis and their clearance by macrophages (Arur et al., 2003), promoting monocyte-to-macrophage differentiation (Yona et al., 2004), and reprogramming their phenotype toward a pro-resolving profile, resulting in modulation of the production of inflammatory mediators.

This modulation was observed through the treatment with CDX-Ac_2-26_ and the reduction of plasma MCPT-1 levels, as well as mast cell degranulation. We know that the DENV activates mast cells and promotes their degranulation (Troupinet al., 2016). Through a study it has been demonstrated that the Ac_2-26_, can inhibit mast cell degranulation mediated via a direct pathway through FPRs (Oliveira et al., 2021), further supporting our findings. AnxA1’s effects depend on its interaction with the FPR2/ALX receptor (Perretti et al., 2009), to which Ac_2-26_ also binds, mimicking the main biological functions of the full-length protein (Takedachi et al., 2025). AnxA1 stands out as a protein known for modulating the inflammatory response and accelerating the return to homeostatic balance in different pathophysiological contexts (Gadipudi et al., 2022; de Araújo et al., 2022).

Notably, CDX-Ac_2-26_ enabled oral administration, representing a considerable advancement in terms of clinical applicability, given the preference for less invasive and more patient-accepted routes (Li et al., 2025). Furthermore, CDX-Ac_2-26_ allowed a three-fold reduction in the intraperitoneal dose, minimizing potential adverse effects and decreasing treatment cost. On the other hand, oral administration required the highest dose, possibly due to increased susceptibility to enzymatic degradation in the gastrointestinal tract (Reinholz et al., 2018), drugs can be inactivated in the gastrointestinal tract, metabolized in the intestinal wall or in the liver before entering systemic circulation, leading to a reduction in the amount that reaches the site of action, this effect is known as first-pass metabolism (McLeod, H. L., 2008). Nevertheless, the therapeutic efficacy observed via the oral route highlights the translational potential of the formulation.

Our *in vivo* evidence reinforces the clinical notion that dengue is a viral-triggers pro-inflammatory disease, in which both the virus-related tissue insult and inflammatory disbalance contribute to mortality. Because all experimental evidence supports that CDX-Ac_2-26_ exhibited an exclusive anti-inflammatory effect and did not interfere with the host’s ability to control the infection, as we detected unchanged viral titers even after treatment. This suggests that viral dissemination plays a crucial role in the pathogenesis of severe dengue, since only 33% of infected mice treated with CDX-Ac_2-26_ survived. This partial protection may reflect an improvement due to inflammation modulation, although it is insufficient for full protection. There are antivirals being tested worldwide for DENV, but without success, some of these include nucleoside analogues such as Sofosbuvir. Sofosbuvir, a nucleotide analogue approved for hepatitis C, has also been tested against several flaviviruses due to its role in inhibiting RNA polymerase, an enzyme essential for the replication of various RNA viruses (Sacramento et al., 2017; de Freitas et al 2019). In our study, Sofosbuvir reduced virus replication and promoted 50% survival. We investigated the potential of combining antiviral and anti-inflammatory therapy. The combination of CDX-Ac_2-26_ and Sofosbuvir promoted 100% survival. We suggest Sofosbuvir inhibits viral replication, while CDX-Ac_2-26_ reduces inflammation.

The therapeutic potential of this and others pro-resolving mediators in treating inflammatory diseases, both infectious (Boff et al., 2020; Lara et al., 2025; Tavares et al., 2022) and sterile (Chen et al., 2023; Woo et al., 2025), has been increasingly explored. Highlighting the therapeutic potential of strategies aimed at restoring endogenous pro-resolving mediators in individuals affected by inflammatory conditions. Our data demonstrated that the CDX-Ac_2-26_ formulation enhanced the therapeutic potential compared to Ac_2-26_. Moreover, it enabled oral administration and was effective at a three-fold lower dose via the intraperitoneal route. Due to its anti-inflammation targeting effect specific, we proposed its combination with an antiviral, which completely prevented DENV-induced lethality. This finding highlights the importance of developing peptide-based formulations that improves its stability and the value of adjuvant strategies targeting severe dengue.

## ABBREVIATIONS

ADE: Antibody-dependent enhancement
AnxA1: Annexin A1
BCRJ: Rio de Janeiro cell bank
CEUA: Animal Ethics Committee
CCL2: Chemokine CCL2
CD: Cyclodextrin
DENV: Dengue virus
ELISA: Enzyme-Linked Immunosorbent Assay
FBS: Bovine serum
FPR2: Formyl Peptide Receptor-2
HP-β-CD: Hydroxypropyl-beta-cyclodextrin
IFN-α/β: Interferon alpha and beta
IFN-γ: Interferon gamma
IL-1β: Interleukin-1β
IL-6: Interleukin-6
IL-8: Interleukin-8
IL-10: Interleukin-10
IRFs: Interferon regulatory factors
KO: Knockout
MCPT-1: Mast cell protease-1
MDA5: Melanoma differentiation associated gene 5
NF-κB: Nuclear factor kappa B
PBS: Phosphate-Buffered Saline
NMR: Nuclear magnetic resonance
RPMI: Roswell Park Memorial Institute
SD: Severe dengue
UFMG: Universidade Federal de Minas Gerais
WT: Wild Type

## AUTHOR CONTRIBUTIONS

J.R.M., V.V.C., M.MT., P.P.G.G., wrote the paper. J.R.M., V.L.B., L.G.R., C.M.Q-J., T.C.M.F., L.S.S., A.S.L.D., A.L.C.S., I.S.L., F.E.O.R., F.R.S.S., J.A.B.P., P.P.G.G., V.V.C., performed the experiments and analyzed data. J.R.M., L.G.R., V.V.C., M.M.T., P.P.G.G., L.P.S., E.S.L., C.M.Q-J., T.M.L.S., provided expertise and contributed to paper writing. M.M.T., P.P.G.G., V.V.C., designed research. All authors have read and agreed to the published version of the manuscript.

## ACKNOWLEDGMENTS

We would like to thank Rosimeire Oliveira, Ilma Marçal, Tania Colina and Hermes Oliveira for their technical assistance. We thank for Women in Science Prize provided by ABC and LOREAL-UNESCO provided to VVC. We also thank CT-Terapias Avançadas e Inovadoras for the support.

## DATA AVAILABILITY STATEMENT

Data and materials may be made available upon written request to the corresponding author.

## DECLARATION OF TRANSPARENCY AND SCIENTIFIC RIGOUR

This Declaration acknowledges that this paper adheres to the principles for transparent reporting and scientific rigour of preclinical research as stated in the BJP guidelines for Design & Analysis and Animal Experimentation and as recommended by funding agencies, publishers and other organizations engaged with supporting research.

---

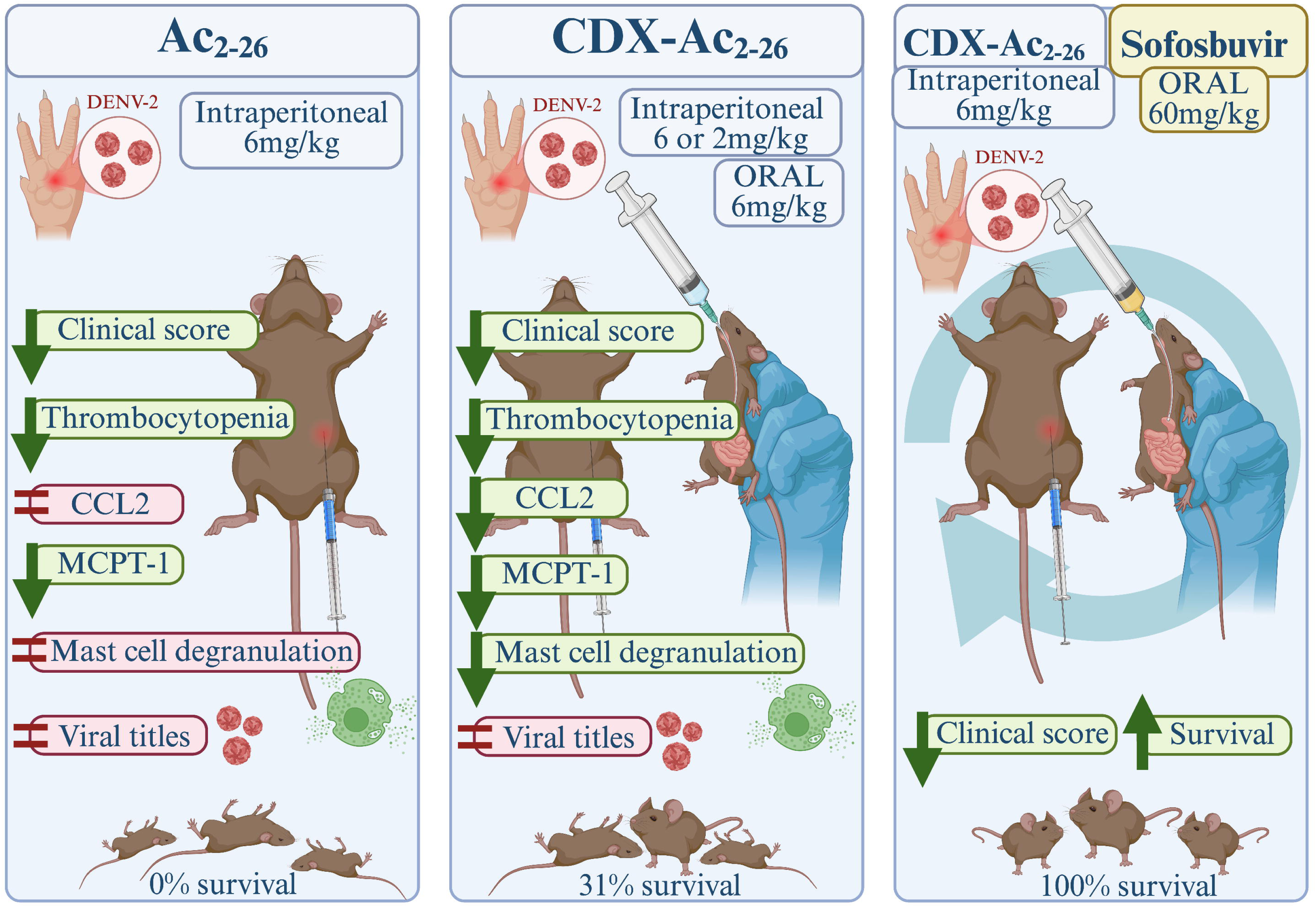

