## Supplementary figures and images for "Cyclodextrin-Based Delivery of the Annexin A1 Mimetic Peptide Ac2-26 Enhances Anti-Inflammatory Effects and Prevents Dengue-Induced Lethality in Combination with Antiviral Therapy"

### Supplementary Figure 1

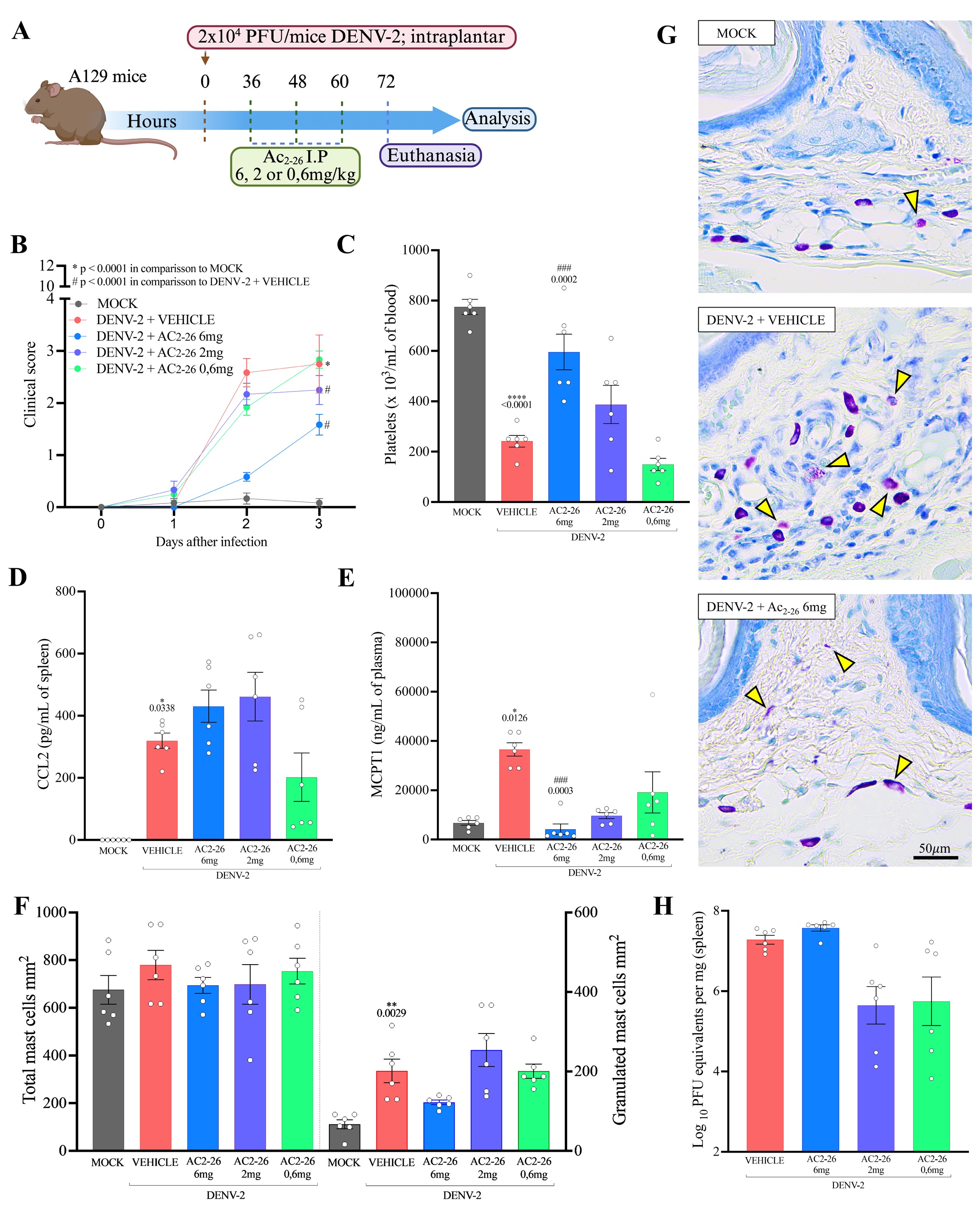

### Supplementary Figure 2

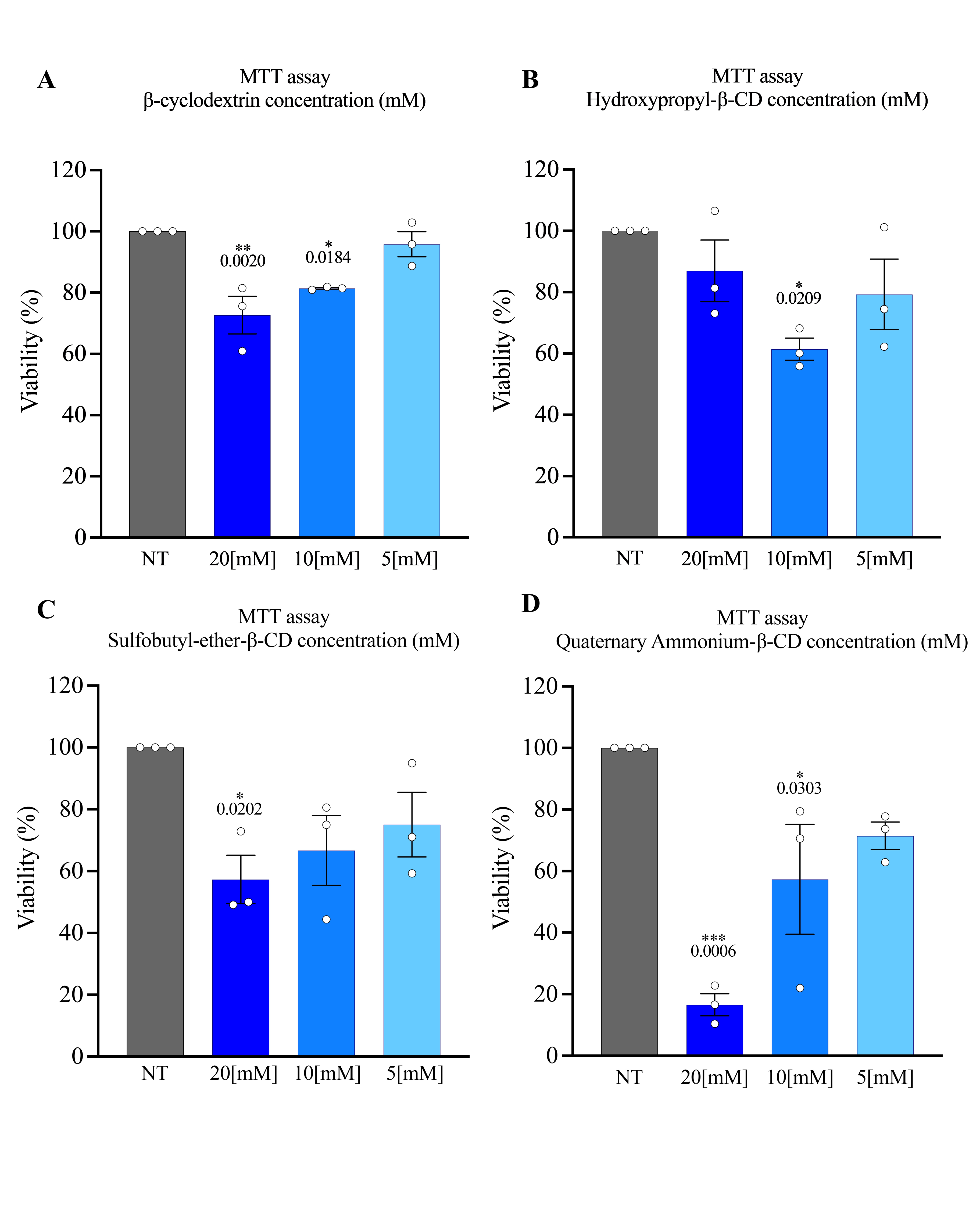
